# Programming conventional electron microscopes for solving ultrahigh-resolution structures of small and macro-molecules

**DOI:** 10.1101/557827

**Authors:** Heng zhou, Feng Luo, Zhipu Luo, Dan Li, Cong Liu, Xueming Li

## Abstract

Microcrystal electron diffraction (MicroED) is becoming a powerful tool in determining the crystal structures of biological macromolecules and small organic compounds. However, wide applications of this technique are still limited by the special requirement for radiation-tolerated movie-mode camera and the lacking of automated data collection method. Herein, we develop a stage-camera synchronization scheme to minimize the hardware requirements and enable the use of the conventional electron cryo-microscope with single-frame CCD camera, which ensures not only the acquisition of ultrahigh-resolution diffraction data but also low cost in practice. This method renders the structure determination of both peptide and small organic compounds at ultrahigh resolution up to ~0.60 Å with unambiguous assignment of nearly all hydrogen atoms. The present work provides a widely applicable solution for routine structure determination of MicroED, and demonstrates the capability of the low-end 120kV microscope with a CCD camera in solving ultra-high resolution structures of both organic compound and biological macromolecules.

## Introduction

Electron diffraction is coming into the scope of the structural biologists and chemists by its capability in determine the structures of small crystals, named MicroED technology^1^. Since 2013, Shi., et al., successfully solved a structure of lysozyme crystal with sub-micrometer size, and demonstrated for the first time that the electron diffraction could be used for the structure determination of small protein crystals. Subsequently, many structures of protein crystals, mostly peptide crystals, were solved using the MicroED method^2^. Moreover, MicroED was recently demonstrated as a powerful approach for determining the crystal structures of small molecular compounds and showed great potential in the related field^3–5^.

Solving the structure of a crystal requires integrated intensities of diffraction spots. Thus, the crystal sample must be rotated continuously with a stable velocity during the data collection such that the Ewald sphere can sweep the entire diffraction spots. Meantime, a fast detector is required to record the diffraction intensities synchronously. This is already a standard design for the modern X-ray diffraction system^6–7^. However, these hardware and software requirements are not always satisfied by commercially-available transmission electron cryo-microscope (cryoTEM). Different strategies have been developed to control the stage (called CompuStage in FEI cryoTEM) and facilitate the data collection, including (1) design of an external device to generate an adjustable constant voltage for rotating CompuStage^8^; (2) use of the FEI TEMSpy interface to set a slow and constant rotation speed^9^; (3) use SerialEM script to control the data collection. For all these strategies, a movie-mode camera with rolling shutter is always required to record images continuously during the stage tilting^10–12^. The direct electron detection (DED) camera also has the movie mode, but is seldom used due to the potential issue of radiation damage to the camera^13^.

While MicroED has shown great potential in determining the structures of small crystals to compensate the traditional X-ray method, most published works using MicroED were still limited to several groups who have the specific camera hardware^1, 3–5,9,14–16^. Meanwhile, most published structures were solved used high-end 200 kV microscopes with field emission gun. In theory, the resolution of electron diffraction is not limited by the contrast transfer function of the objective lens. And hence, low-end electron microscopes, such as the 120 kV microscope with LaB6 gun, should also be able to achieve atomic resolution under diffraction mode. Considering the 200-kV and 120-kV electron microscopes widely installed, removing the requirement for an expensive movie-mode camera and even realizing MicroED on the low-end microscopes is of great importance to boost wider applications of MicroED technology. Meantime, what’s the resolution limit for the current cryoTEM instruments is also a very important information to explore.

Here, we developed a scheme of stage-camera synchronization for cryoTEM and implemented it in a software named eTasED to facilitate the MicroED data collection without the requirement for movie-mode camera and any modification to the current cryoTEM system, which presumably can be used on any cryoTEM with any cameras. Benefiting from eTasED, we collected high-quality data on an FEI Tecnai F20 200-kV microscope (F20) with a Gatan US4000 CCD camera (US4000) and an FEI Tecnai Spirit 120kV microscope (T12) with an FEI Eagle CCD camera (Eagle). Moreover, the applications of our method on several testing samples, a peptide (FUS LC RAC1) and two small organic compounds (acetaminophen and biotin), demonstrated the feasibility of determination of structure at ultrahigh-resolution up to ~0.6 Å with assignment of nearly all of the hydrogen atoms in the structures. The comparisons among the determined structures provided deep insights to not only the behavior of electron diffraction at ultrahigh resolution range but also the advantage of the electron diffraction in resolving the hydrogen atoms which is essential for both biological sciences and drug discovery.

## Result

### Stage-camera synchronization for conventional electron cryo-microscope

To record the diffractions from a rotating crystal continuously, the camera must be exposed continuously and be able to output image frames at a fixed frame-rate^17^. More importantly, no interval exists between adjacent image frames to ensure all diffraction signals are recorded. This is how a movie-mode camera operates. However, for a nonmovie-mode camera, the exposure is not continuous. Each exposure can only output one single frame, and typically requires three steps to finish an exposure cycle. In the first step, the camera is initialized, typically requiring hundreds of milliseconds or several seconds. In this step, the shutter is closed to prevent the sample from being illuminated. In the second step, the camera shutter is turned on to allow for the electron beam to penetrate the sample and reach the camera; simultaneously, the camera begins to record an image until reaching a given exposure time. After the exposure is completed, the shutter is closed. The third step is to readout the image and output the image file to a storage device, which typically requires many seconds depending on the image size and output speed. The total time spending on such an exposure cycle is typically much longer than the given exposure time, i.e., only the second step is the effective exposure. To record the diffraction intensities without missing any diffraction signals, the crystal or the stage must stop tilting within the first and third steps, and the beam should be blanked when the tilting is stopped to avoid unnecessary radiation damage to the crystal.

We used a timer to synchronize the stage tilting with the camera exposure (Fig. 1). In the beginning, a request for exposure is sent to the camera to trigger an exposure cycle, and the timer is turned on simultaneously. The timer will count the elapsed time until reaching the timeline when the initialization of the camera is complete; subsequently, a request is sent to the stage to trigger a continuous tilting for a given angle range. The tilting speed must be measured precisely in order to calculate the elapsed time of tilting before the data collection such that the stage tilting and effective exposure can be started and stopped at the same time. Further, the exposure time must be set to this tilting time. Such setting is to ensure that the exposure is just finished at the same time that the stage reaches the given tilting angle. After the exposure is finished, the entire system will wait for the camera to finish the readout and return to the ready state. This tilting-exposure cycle will be performed repeatedly until reaching the final tilting angle.

**Figure 1.**
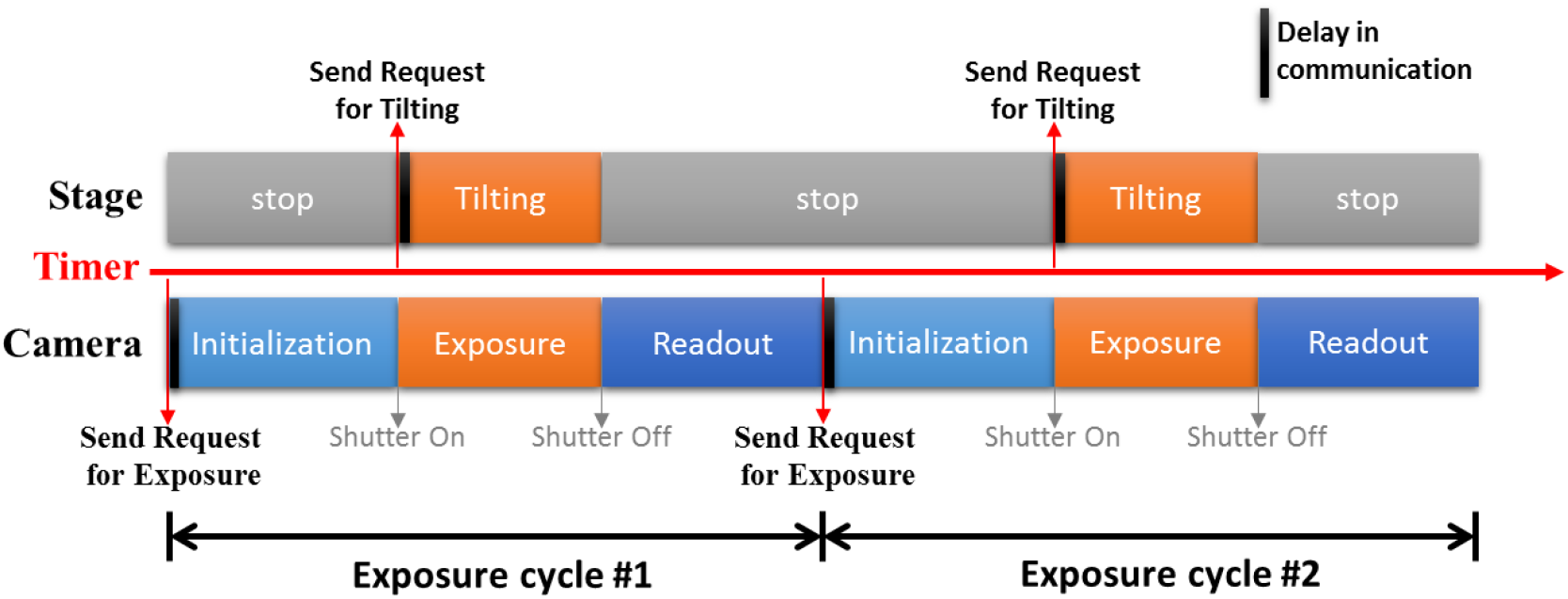
Schematic diagram of the stage-camera synchronization. The two colorful bars indicate the operations of the stage and camera along the time line, respectively. The horizontal red arrow denotes the timer that controls the time points (the vertical red arrows) to send requests for operations. After the request is sent out, the camera or the stage requires some time to receive the request and generate a response, which causes a delay (the vertical back line) in the synchronization. Two exposure cycles are shown.

Two key parameters used in the synchronization are the tilting speed of the stage and the initialization time of the camera. The tilting speed can be measured easily by averaging the speed of rotating a large angle. The initialization time of the camera varies with different cameras, but is usually a constant for a specific camera under specific settings. For example, the initialization time of our US4000 on F20 is 520 ms, and that of our Eagle on T12 is 1625 ms. A protocol has been developed to estimate the camera initialization time (see Methods).

The stability of the stage during continuous tilting is important for the data quality. Uneven or unstable tilting results in errors in the recorded diffraction intensities. To assess the stage ability in our synchronization scheme, a Kikuchi line method was developed to trace the tilting of the stage under the condition of MicroED data collection (see Methods, Supplementary Figure 1 and 2). We found that the CompuStage in our F20 is sufficiently stable at all tilting angles, and the influences of the acceleration and deceleration was negligible under an exposure time as long as several seconds.

This synchronization scheme is also available for a movie-mode camera. With a movie-mode camera, the exposure time can be set much longer than that of a single-frame camera, such that each tilting-exposure cycle can cover a large or even the entire range of the tilting angle. Meanwhile, it typically takes many minutes or more than a hundred frames to collect a set of data for a single crystal. Some movie-mode cameras limit the total exposure time and the total number of frames. This synchronization scheme also removes the requirement for a long exposure time of many minutes, and hence helps to avoid such limitation.

### Ultrahigh-resolution diffractions of peptide nanocrystals from 200- and 120-kV electron microscopes

The synchronization method was tested on the two cryoTEMs mentioned above, F20 and T12, using a peptide crystal (FUS LC RAC1) reported in our previous work^16^. F20 is equipped with a field-emission gun and a US4000 camera, and T12 has a LaB6 gun and an Eagle camera. A relative titling speed of 0.6% were set for both microscopes, and led to similar tilting speeds, 0.1748 ^o^/s on F20 and 0.1736 ^o^/s on T12. The tilting angle range for each exposure was set to 1°, accordingly, the exposure time was 5.72s for US4000 and 5.76s for Eagle. For each microscope, a dataset from the best single crystal and a dataset merged from multiple crystals were used to demonstrate the data quality. Most crystals observed on the two microscopes can diffract to ~0.6 Å resolution. The best ones can be exposed and tilted for more than 90°, corresponding to a data completeness of ~60% under the *P21* space group, and diffract up to 0.58 Å and 0.57 Å resolution on F20 and T12 (Fig. 2), respectively.

**Figure 2.**
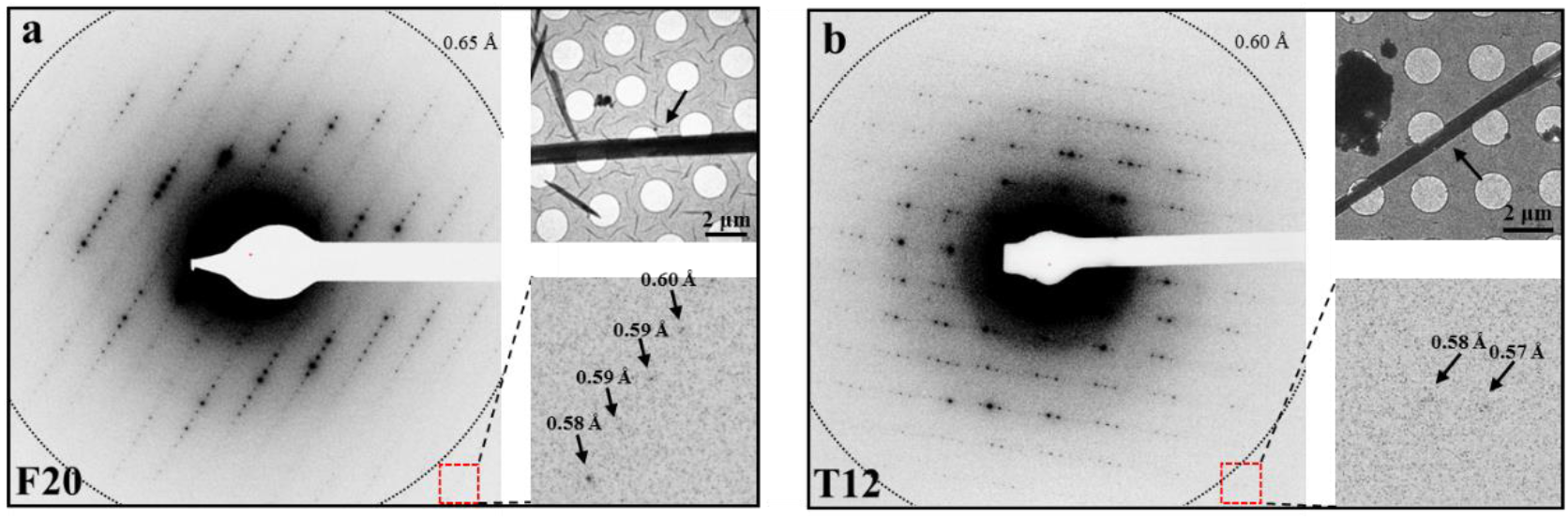
Typical diffraction patterns of FUS LC RAC1 crystal. **a)** and **b)** are diffraction patterns collected on F20 and T12, respectively. The red crosses in the center of the diffraction patters indicate the beam center, and the dotted rings indicate the final resolution of the solved structures. The arrows on the magnified images of the red boxes indicate some high-resolution diffractions. The corresponding images of the needle-like crystals (pointed by black arrows) are shown on the top right of each panel.

The overall data quality was first evaluated based on the data processing. All the datasets from F20 were processed successfully by XDS^18^ with *R*_merge_ between 0.08 and 0.15. The initial overall *R*_merge_ of each single-crystal T12 datasets was high in the beginning of the data processing using XDS. Some of them reached beyond 0.5. After optimizing the data quality in XDS by refining the parameters of beam divergence, reflecting range and their deviations, the overall *R*_merge_ dropped to ~0.2. The crystal mosaicity of the optimized T12 datasets was more than 1°, which is higher than that of the F20 datasets, primarily less than 0.3°. Accordingly, a REFLECTING_RANGE parameter defined in XDS reached 6~8° for the T12 datasets, which meant it took 6~8° of rotation for a reflection to pass completely through the Ewald sphere on the shortest route. Not only the overall *R*_merge_ but also the overall <l/σ(I)> of the F20 datasets are better than those of the T12 datasets (Supplementary Table 1). These effects may relate to some systematic inaccuracy of the T12 dataset, which is reasonable if considering the lower optical performance of T12 than that of F20.

The statistics in different resolution shells were further compared (Fig. 3). The completeness of the merged F20 dataset is slightly higher than that of the merged T12 dataset from the lowest resolution to ~0.75 Å. Beyond ~0.7 Å to a higher resolution, the T12 dataset indicates higher completeness than the corresponding F20 datasets (Fig. 3a), implying that more high-resolution reflections are recorded on T12. The <*I/σ(I)*> of the merged F20 dataset in the range from the lowest resolution to ~0.85 Å are much higher than that of the merged T12 dataset but become lower in the range beyond ~0.8 Å (Fig. 3b and Supplementary Table 1a and 1b). The trends from the completeness and σ*I/σ(I)*> imply that the merged T12 dataset have better signal-to-noise ratio (SNR) and more reflections in the high-resolution range beyond ~0.8 Å. In the resolution range lower than ~0.8 Å, the *R*_merge_ of the merged F20 dataset is better than that of the merged T12 dataset, and becomes worse at higher resolution range (Fig. 3c). For CC_1/2_^19^ (the correlation coefficients between randomly divided half datasets) the merged F20 datasets always exhibited better quality than the merged T12 datasets at all resolution shells (Fig. 3d). Similar trends for the merged datasets were also observed for the single-crystal datasets (Fig. 3). Combining all the four statistics parameters above, it is clearly seen that the 120-kV T12 yield stronger (or high SNR) and more detectable reflections at resolution range higher than ~0.8 Å resolution, but which have worse accuracy. The reason is still not clear. One possibility is that the lower electron energy (from 120-kV T12) leads to stronger interactions between the electrons and materials, and hence stronger and more diffractions at high-resolution range, but the stronger interactions may also lead to stronger multiple scattering and reduce the repetitiveness of the reflections, especially, the high-resolution reflections.

**Figure 3.**
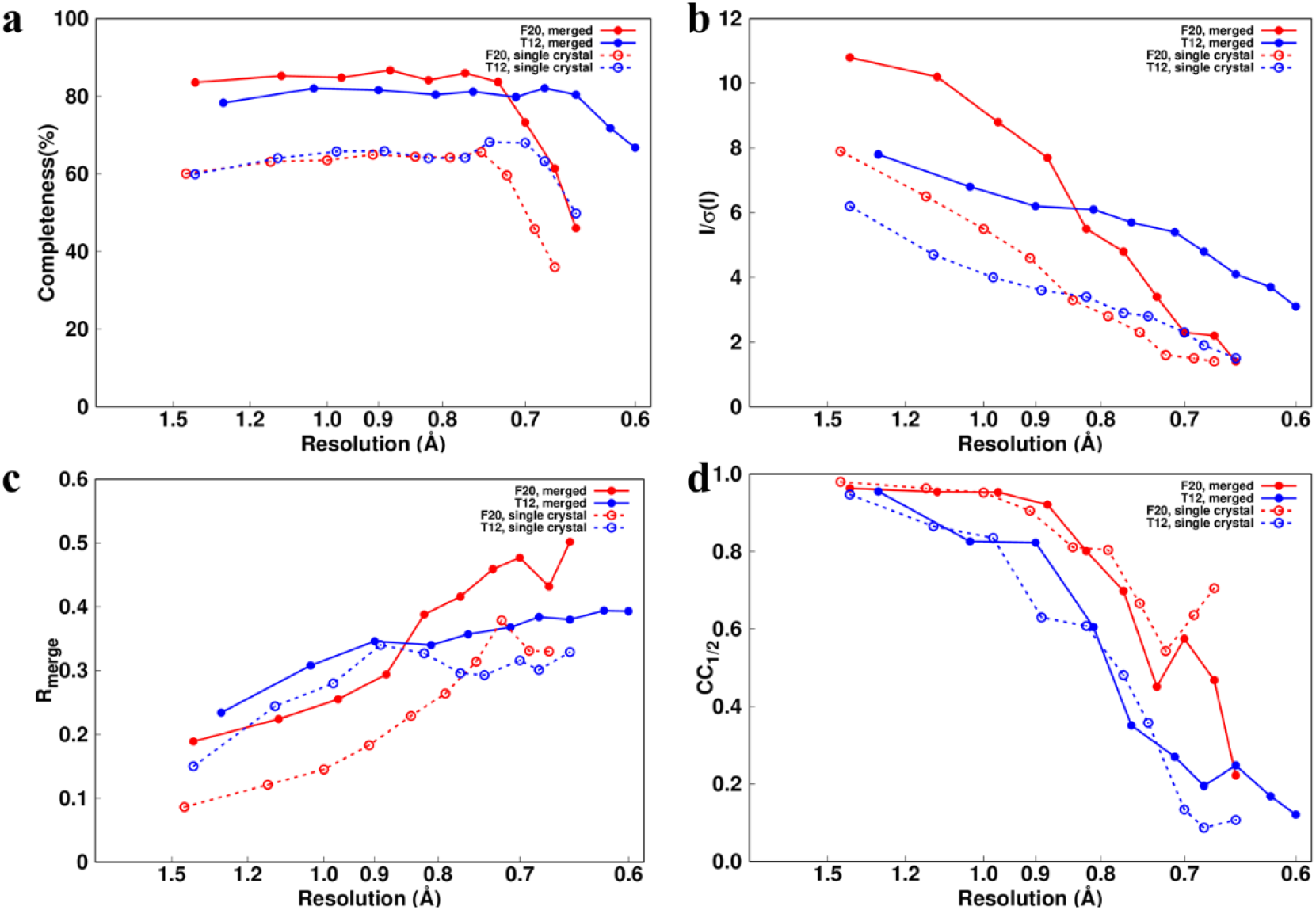
Statistics of the diffraction data of FUS LC RAC1 crystal. **a), b), c)** and **d)** are the curves of the completeness, I/σ(I), R_merge_ and CC_1/2_ against the resolution, respectively. These curves were calculated from the merged (red solid lines) and single-crystal (red dashed lines) F20 datasets, and the merged (blue solid lines) and single-crystal (blue dashed lines) T12 datasets.

Although performing worse than F20 in terms of overall quality, T12 still yielded high-quality diffractions sufficient to solve a ultrahigh-resolution structure (discussed later). The final resolutions of the determined structures of the merged datasets from F20 and T12 were cut-off at 0.65 Å and 0.60 Å (Fig.4a and b, Supplementary Table 2), respectively, and those of the single-crystal datasets from F20 and T12 were cut-off at 0.67 Å and 0.65 Å, respectively, based on the threshold of <*I/σ(I)*> ≈ 1.5. The data quality was further compared with all the published structures (all used movie-mode cameras) solved by the MicroED method before 2019 (Supplementary Table 3). The data statistics of <*I/σ*(*I*)> from the merged datasets are better than those of most published work. Particularly, the current datasets from both the F20 and T12 exhibit the highest resolution by MicroED. The averaged B-factor (MeanB, the last column in Supplementary Table 3) of our refined models are much lower than most of the published structures solved by MicroED, which demonstrates the high data quality obtained using eTasED.

**Figure 4.**
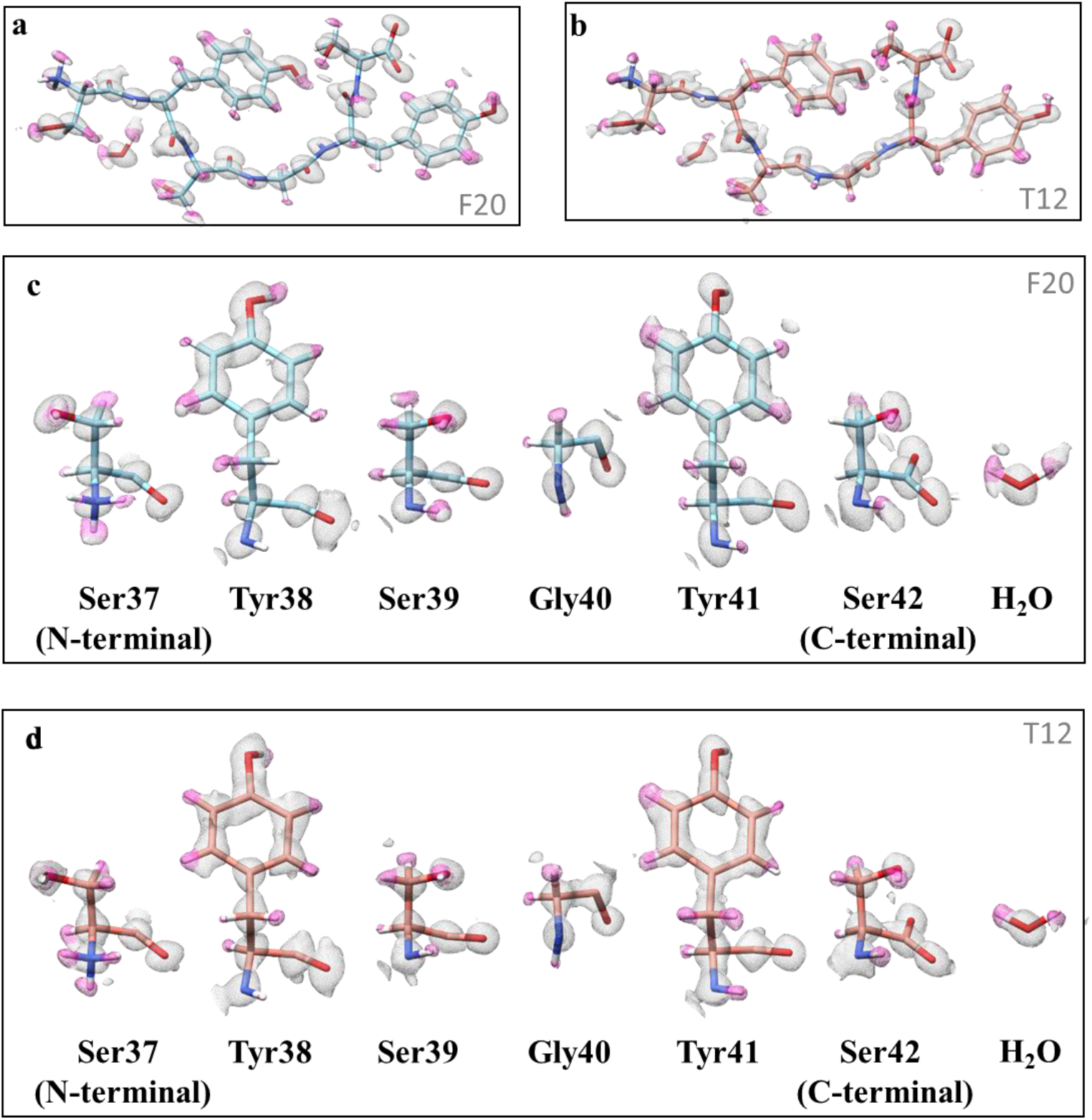
Structures solved from the merged datasets. **a)** and **b)** are 2F_o_-F_c_ density maps solved from the merged datasets from F20 and T12, respectively. **c)** and **d)** are the magnified density maps of the residues in **a)** and **b)**, respectively. The densities near the hydrogen atoms are colored in pink. The contour thresholds of all the maps are at 1.0 σ.

### Determination and analysis of the peptide structure at ultrahigh resolution

The diffractions up to ~0.6 Å resolution enabled the direct method for phasing the diffractions from the peptide crystal mentioned above (Supplementary Figure 3 and 4). The two merged datasets from F20 and T12 yielded correct solutions by SHELXD^20^ (Supplementary Figure 3a and b). More than 10 single-crystal datasets collected on F20 reached the completeness of ~60% in the resolution shells ranging from 1.5 Å to 0.7 Å, and exhibited the overall *R*_merge_ of less than 15%, which enabled the use of the direct method for each single crystal individually (Supplementary Figure 3c). Whereas, the same attempts for all the single-crystal datasets from T12 failed, and did not yield the correct structures by the direct method even after extensive trials with SHELXD (Supplementary Figure 3d). Even though the single-crystal dataset from T12 exhibited a similar unique reflection number (~3,700), multiplicity (~2.5), and slightly better overall completeness compared to those from F20 (Supplementary Table 1c and 1d). The noise level of the single-crystal T12 dataset, indicated by the distribution of *σ(I)*, are higher than that of the two F20 datasets as well as the merged T12 dataset (Supplementary Figure 5), which results in low accuracy for the measured intensities, especially, for the intensities of the relatively weak reflections. These comparisons together with the discussions mentioned before comprehensively indicate that the measured diffraction intensities from T12 are less accurate at high resolution than those from F20, which might explain why the single-crystal T12 dataset did not produce the correct structure. However, the accuracy of the T12 dataset can be significantly improved after merging the intensities from multiple crystals, demonstrated by the success of the direct method for the merged T12 dataset (Supplementary Figure 3b). Molecular replacement was subsequently used for solving the structure of the single-crystal dataset from T12 and produced the correct solution (Supplementary Figure 4d).

The ultrahigh-resolution structure of FUS LC RAC1 peptide reveals an ordered-coil amyloid spine. Almost all of the hydrogen atoms are clearly visible at the *2Fo-Fc* map (the contour level at 1.0σ), including the two hydrogen atoms of the water molecule and three hydrogen atoms of the protonated amino group on Ser37 (Figure 4). The hydrogen atoms are also clearly revealed using the single-crystal datasets from F20 and T12 (Supplementary Figure 6). Unambiguous assignment of the accurate position of each hydrogen enables us to build up a precise and complete hydrogen-bond network formed by intra-molecular as well as inter-molecular interaction in the RAC1 peptide structure (Supplementary Figure 7). The architecture of the RAC1 peptide in crystal lattice recapitulates the self-assembled structure of RAC1 in amyloid fibril, where the hydrogen-bond network is regarded as a main force to maintain the fibril structure and regulate its dynamic assembly^16^. Therefore, the unambiguous assignment of the entire hydrogen-bond network from the ~0.6 Å peptide structure may provide valuable resource for accurate calculation of the geometry and energy of hydrogen-bond network in stabilizing the amyloid fibril spine.

### Ultrahigh-resolution structural determination of two organic compounds

Because of the recent successful applications of MicroED on small organic compounds^3–5^, we examined eTasED programmed electron microscope in determining the structures of two organic compounds including acetaminophen (Aladdin Company) and biotin (Thermo Fisher Scientific Inc.) from commercial available powder, using our 200-kV F20 and 120-kV T12 with CCD cameras mentioned above. Small amount the powder was grounded, deposited on the grid, frozen in liquid nitrogen, and transferred to the microscopes. By using eTasED, we collected ultrahigh-resolution diffraction data of both compounds with high quality (Supplementary Table 4). The best data were achieved for acetaminophen which diffracted up to 0.54 Å and 0.51 Å on F20 and T12, respectively (Supplementary Figure 8a and b). Regardless the data accuracy influenced by the multiple scattering and optical aberrations, this results demonstrated the capability of low-end T12 in delivering the detectable diffraction information up to 0.51 Å resolution. The corresponding structures were determined at 0.65 Å and 0.83 Å resolutions, respectively (Fig. 5a and b). Benefiting from the ultrahigh-resolutions, most of the hydrogen atoms are clearly visible in both the 2Fo-Fc maps and hydrogen atom omitted Fo-Fc maps calculated by SHELXL, which demonstrated the capability of the 120-kV T12 with CCD camera as well as our data accusation scheme. Unlike acetaminophen, the diffractions from biotin crystal can only reach up to 0.68 Å and 0.84 Å resolution on F20 and T12, respectively (Supplementary Figure 8c and d). And the corresponding structures were determined at 0.75 Å and 0.90 Å resolutions, respectively (Fig. 5c and d). Accordingly, the hydrogen atoms are not well defined in the resolved maps of biotin.

**Figure 5.**
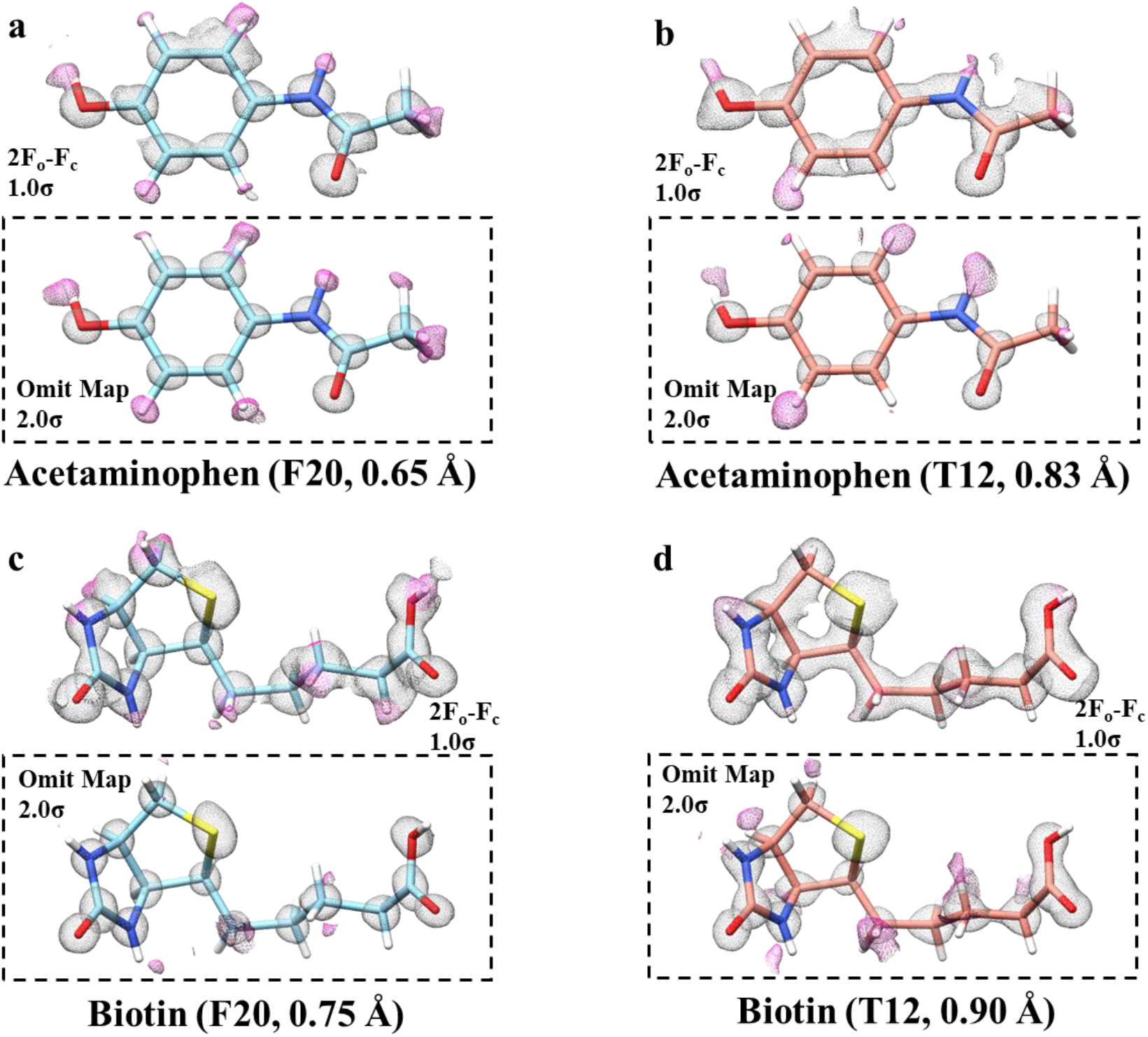
Structures and density maps of acetaminophen and biotin. **a)** and **b)** are 2F_o_-F_c_ density maps and omit maps of acetaminophen from F20 and T12, respectively. **c)** and **d)** are 2F_o_-F_c_ density maps and omit maps of biotin from F20 and T12, respectively. In 2F_o_-F_c_ maps, the densities near the hydrogen atoms are colored in pink, and the contour thresholds were set to 1.0 σ. For the maps in the dashed box, the gray densities are the 2F_o_-F_c_ maps with the contour thresholds set to 2.0σ, and the pink densities are the omit Fo-Fc map with the contour thresholds set to 2.0σ. The name of cryoTEMs and resolutions are labeled in the bottom of each panel.

Taken together, these results demonstrated that both the 200-kV and 120-kV cryoTEM with CCD cameras are able to generate ultrahigh-resolution data for the small crystals of organic compounds. The quality of the diffraction data and the resolution of the determined structure of the two compounds are comparable or even much better than those obtained by advanced Talos Arctica cryoTEM with a new-released CetaD camera^4^. More importantly, our result demonstrates that the low-end 120kV electron microscope and CCD cameras with eTasED is sufficient to determine the high resolution structures of small organic compound, and removes the high barriers of the hardware requirements for applying MicroED in the field of chemistry especially organic, medicinal chemistry, and material science.

## Discussion

In this study, by synchronizing the stage and the camera, we developed an automated scheme to collect the diffraction data for MicroED. This method allows for the use of any type of camera, including the non-movie-mode camera, to record the diffraction intensities from a continuously tilted crystal. The best diffraction recorded using the 120-kV T12 with CCD camera in the present work is at 0.51 Å resolution, which implies the capability of these low-end instrument. Such feature significantly reduces the hardware obstacle for most users, and makes widely equipped CCD cameras installed on 200-kV or 120-kV electron microscope immediately available for MicroED. Therefore, nearly any electron cryo-microscopes can be used for MicroED without changes to their hardware and software. The present work will boost the applications of MicroED in more cryoEM laboratories for studying both biological macromolecules and small compounds.

The 120-kV T12 was proven to be available for MicroED and could achieve a high resolution of up to ~0.60 Å in the present work. Of note, the data quality from the 120-kV T12 was slightly worse than that from the 200-kV F20. However, our T12 could still reveal the hydrogen atoms with the similar quality as F20. This implies that the expenses for MicroED can be reduced significantly using a low-end electron microscope with a CCD camera. It is also noteworthy that the performance of the camera is always a key factor for the data quality, especially, the signal-to-noise ratio (SNR) for the weak signal. Both US4000 and Eagle had been developed as main-stream detectors for imaging before the breakthrough of DED camera, and hence, as shown in the present work, their SNR is sufficient for recording the diffraction intensities.

Until now all the published MicroED works use the CompuStage (FEI company) to tilting the sample. However, the behavior of the CompuStage under continuous tilting was never examined. This brings potential risk in evaluating the errors in the collected data. We examined the CompuStage on our F20 using two tracking markers: gold nanoparticles observed at the image mode and Kikuchi patterns observed at the diffraction mode. Because the diffraction is invariant to the sample translation, the Kikuchi patterns provide an accurate method to observe the tilting stability of the CompuStage. Under a low speed, ~0.2 °/s, the tilting of the CompuStage is smooth and stable. The acceleration and deceleration of the CompuStage were observed in the beginning and end of the tilting, respectively. However, their influence to the data quality should be minor when the exposure time is long. The influence becomes significant only when an extremely short exposure time is used. This should be avoided in the application of the present method.

The synchronization scheme for MicroED has been implemented in the eTasED program, which is a Microsoft™ Windows program written in Visual C++. It contains three user-friendly graphic interfaces, including a camera server program to control the camera, a TEM server to control the electron microscope through FEI scripting interface, and a main program for data collection. The current version of the program can be installed on an FEI electron microscope with most mainstream cameras. The program is available for downloading through http://github.com/THUEM/eTasED.

## Acknowledgements

This work was supported by funds from The National Key Research and Development Program (2016YFA0501902 and 2016YFA0501102 to X.L. and C.L.), National Natural Science Foundation of China (31570730 and 31722015 to X.L. 91853113 and 31872716 to C.L.), Advanced Innovation Center for Structural Biology (to X. L.), Tsinghua-Peking Joint Center for Life Sciences (to X. L.), and One-Thousand Talent Program by the State Council of China (to X. L. and C.L.). We acknowledge Tsinghua University Branch of China National Center for Protein Sciences Beijing for providing facility support.

## Contributions

X.L. and C.L. initialized the project. X.L. designed the method and wrote the program. H.Z. and F.L performed all the experiments, data collections and TEM observations. F.L. and D.L. prepared and optimized the crystal. Z.L. and H.Z. analyzed the diffraction data and solved the structures. X.L. wrote the manuscript. All authors contributed to the data analysis and manuscript revision.

## Code availability

The package of eTasED and documents for noncommercial use is available at http://github.com/THUEM/eTasED. A protocol and manual will be included in the package.

## Data availability and Accession Code Availability Statements

The models from the merged and single-crystal datasets from F20 and T12 are deposited into Protein Data Band with entry codes XXXX, XXXX, XXXX, XXXX, XXXX, and XXXX, respectively. The raw images support the findings of this study are available from the corresponding author upon request.

## Methods

### Crystallization

FUS LC RAC1 peptides were synthesized by ChinaPeptides. Peptides were dissolved at a 20 mg/ml concentration in ddH2O. Droplets of FUS LC RAC1 solution were mixed in a 1:1 ratio with 1.95 M ammonium citrate, pH 7.0, by hanging-drop vaporization. Crystals were grown at 16 °C.

Acetaminophen powder was purchased from Aladdin Company. Biotin powder was purchased from Thermo Fisher Scientific Inc.

### Sample Preparation

The FUS LC RAC1 crystals were clumped heavily together. We used a microtool from Hampton to separate the clumps; subsequently, the samples were diluted four-fold using the crystallization buffer. A drop (4 μL) of suspended crystals was loaded on one side of glow-discharged Quantifoil EM grids (R2/2, Cu 300 mesh; QUANTIFOIL Company). The grid was then blotted from the other side and washed twice manually with 4 μL 5% v/v PEG 200 buffer. Finally, the grid was blotted twice and vitrified by plunging into liquid ethane using a Vitrobot Mark IV (FEI Company). The frozen grids were transferred to F20 or T12 using a Gatan 626 cryo-holder.

Small amount of acetaminophen or biotin powder was ground using a mortar and transferred to a glass slide. We then used a glow-discharged Quantifoil EM grids (R2/1, Cu 300 mesh; QUANTIFOIL Company) to gently touch the powder for several times. Then the grid was gently shook to remove excessive powder, and then plunged into liquid nitrogen and transferred to F20 or T12 using a Gatan 626 cryo-holder.

### Stability measurement of the CompuStage during continuous tilting

To optimize the control of the stage and determine the potential problems influencing the data quality, we examined the single-tilt stage on our F20 under both the image mode and diffraction mode. Under the image mode, we used the gold nanoparticles distributed on continuous carbon film to track the movement of the CompuStage (Supplementary Figure 1a, Supplementary Movie 1). By positioning the sample slightly off the eucentric height, stage tilting was presented by the translation of the particles along the direction of the tilting. Meantime, the movement of the nanoparticles is also sensitive to the translational instability of the CompuStage during the tilting. Under the diffraction mode, we used the Kikuchi patterns of a thin silicon crystal to measure the stage rotation (Supplementary Figure 2a and b, Supplementary Movie 2). The Kikuchi pattern is invariant of the translation^21^, i.e., it is free of the influence of any stage translations. Therefore, the Kikuchi pattern is a good indicator reflecting the tilting angle. A DE20 camera (DE company) operating at the movie mode was used to record the sample tilting under both the image and diffraction mode with a frame rate of 25 frames/s.

Three interfaces control the stage tilt for an FEI electron microscope: the push button on the control panel, the TEMSpy interface, and the scripting interface. The first two have been used in several published works, but are not suitable for the programmed control. We chose the scripting interface and used software to control the rotation. The stage can be tilted using one of two methods through scripting. One is without speed control and uses only the default speed. For this method, the tilting speed is fast. We attempted to decelerate the tilting by sending requests of small-angle tilts sequentially (0.01° per tilt). This method has been used to solve the structure of the FUS LC RAC1 peptide crystal and achieved a 0.73 Å resolution^16^. However, the speed measurement by Kikuchi patterns and nanoparticles demonstrated highly uneven tilting speed and strong stage shaking in translation. Another method is with speed control. The nominal speed for the FEI CompuStage is 29.7°/s in 100% speed. The speed can be controlled by changing the percentage parameter. This speed control is the same as that used by TEMSpy. Considering the problems in the first method, we used the second method in this work and implemented it in eTasED.

We tested the tilting of 1.4° with 1% speed. Under the image mode, the stage appeared highly unstable, and exhibited strong shaking along the direction of the tilting (Supplementary Figure 1, Supplementary Movie 1). Under the diffraction mode, the Kikuchi lines moves smoothly (Supplementary Movie 2). By measuring the movement of a cross point of two Kikuchi lines, we traced and calculated the relative tilting angles. We observed that the tilting was smooth (Supplementary Figure 2c). Therefore, the translation instability did not influence the stability of the tilting. A concern is the acceleration and deceleration observed at the beginning and the end of the tilting, respectively. Fortunately, both of them are moderate and take only a short time of ~40 ms (Supplementary Figure 2d). The errors caused by the acceleration and deceleration could be negligible when the exposure time is much longer than 40 ms, for example, more than 5 s in the present work. The tilting angle and speed (1% speed) with the given tilting range of 1.4° in different starting angles from −60° to 60° were also tested (Supplementary Figure 2e and f). The speed is even and does not dependent on the staring angle. Unexpectedly, the measured tilting angles (Supplementary Figure 2e) is always slightly less than the nominal tilting angle of 1.4°, corresponding to ~0.01° errors (~0.7% of 1.4°). These errors may cause inaccuracy or problems in the measurement of the diffraction intensities.

### Camera initialization time test

The camera initialization time was measured by a build-in module in eTasED. When starting the camera initial time measurement in eTasED, a series of exposure cycles would be performed automatically. In each cycle, the beam was turned on, and a request to trigger the exposure was sent to the camera, then the beam was blanked using the beam blanker provided in FEI scripting interface after a short time interval, and the averaged pixel intensity was read from the camera after the exposure finished. This time interval was increased by a small step (a given value by the user), and the procedure above would be repeated continually until the time interval was longer than the time to finish an exposure cycle. A curve of the averaged pixel intensity versus the time interval would be shown on the interface of eTasED. If the time interval was shorter than the camera initial time, the camera would not be exposure, accordingly, the averaged pixel intensity would be zero. Otherwise, it would exhibit a non-zero value. The inflection point of the curve was regarded as the camera initialization time. The precision of the measurement can be adjusted by changing the time interval, typically, 0.05 s.

### Data collection

The 200-kV diffraction data and images of crystals were observed using F20 and recorded using the US4000 camera with a sensor size of 4,096 × 4,096 pixels. The eTasED software was used to perform the semi-automated MicroED data collection. The nominal camera length was set to 520 mm and calibrated by an oriented-gold-film standard sample. The exposure time for each image frame was 5.72 s. The selected area aperture (diameter of 200 μm, equivalent to ~5 μm on the object plane) was used to select area for diffraction. The measured electron dose on the sample was approximately 0.01 e^-^/(Å^2^×s). During the exposure time, crystals were rotated continuously at a speed of 0.1748°/s per second. Therefore, each image frame covered the 1.0° wedge, and each dataset from a single crystal covered the angle range from 50° to 101° in the reciprocal space. The best single crystal data set of FUS LC RAC1 from F20 was obtained by rotating the crystal from −40° to 49° (Supplementary Table 1c).

The 120-kV diffraction data and images of crystals were observed using T12 and recorded using the Eagle camera with a sensor size of 4,096×4,096 pixels. The eTasED software was used to perform the semi-automated MicroED data collection. The nominal camera length was set to 500 mm and calibrated by a multi-crystal aluminum standard sample. The exposure time for each image frame was 5.76 s. The selected area aperture (diameter of 200 μm, equivalent to ~5.5μm on object plane) was used to limit the sample area for diffraction. The measured electron dose on the sample was approximately 0.01 e^-^/(Å^2^×s). During the exposure time, the crystals were rotated continuously at a speed of 0.1736°/s per second. Therefore, each image frame covered the 1.0° wedge, and each dataset from a single crystal covered the total 40° to 113° angle range in the reciprocal space. The best single crystal data set of FUS LC RAC1 from T12 was obtained by rotating the crystal from −65° to 48° (Supplementary Table 1d).

### Data processing and structure determination

Diffraction images were collected as the MRC format and converted to the SMV crystallographic format using cryoec_diffview in the eTasED suite. The electron diffraction data were processed by XDSGUI in XDS package. The beam center of each dataset was obtained by the cryoec_diffview software and loaded into XDS. To search for the strong diffractions, the parameters of STRONG_PIXEL and MINIMUM_NUMBER_OF_PIXELS_IN_A_SPOT in XDS were set to 10.0 and 4.0, respectively, and TEST_RESOLUTION_RANGE was set from 5.00 Å to 1.00 Å for indexing owing to the lack of low resolution data. These settings are critical for indexing all the datasets correctly. Owing to the high *R*_merge_ of the single-crystal T12 dataset, further data quality optimizing was performed for this dataset in XDSGUI, then the datasets were re-integrated. Finally, the most isomorphous data, analyzed by xds_nonisomorphism, were merged together with XSCALE in XDS package and then converted to mtz files with XDSCONV in XDS package. The statistics of data collection were obtained by phenix.merging_statistics^22^ and listed in Supplementary Table 1. XPREP in SHELXTL package was used to prepare the input files for the direct method phasing using SHELXD. The electron density map was calculated by SHLEXL in SHELX package. Coot^23^ was used to build the amino acid manually into the electron density map to match with the model given by SHLEXD. The single-crystal dataset recorded on T12 was solved by molecular replacement using PHASER^24^ supplied with the refined model of the merged T12 dataset. Crystallographic refinements were performed using REFMAC5^25^ with electron atomic scattering factors with the keyword file containing “source EM” Data processing and refinement statistics are listed in Supplementary Table 2. All-atom clashscores of the two structures by MolProbity^26^ statistics are both 0.00. The Ramachandran plot indicated 0% outliers and 100% favored.

The electron diffraction data of acetaminophen and biotin were processed the same way as above. The structures were solved by SHELXD and refined by SHLEXL with proper geometrical restrains using electron atomic structure factor^27–28^. The hydrogen atoms were added into the structures using HFIX and validated by hydrogen atoms omitted Fo-Fc map calculated by SHLEXL.

### Kikuchi pattern data collection and processing

A thin single-crystal silicon was used to produce the Kikuchi pattern. The silicon sample was transferred to microscope using a Gatan 626 cryo-holder but without liquid nitrogen cooling. A DE20 camera operating at the movie mode was used to record the sample tilting with a frame rate of 25 frames/s. The values of the camera initial time and exposure time in eTasED were set slightly larger than the true values to ensure that the movie contained the whole process of a tilt. Then a movie of Kikuchi pattern moving with stage tilting for 1.4° with 1% speed was collected. The same procedure was repeated at different starting angles from −60° to 60° to investigate the stability of the stage in different angles. These movies were processed using a custom-made MATLAB script, measuring the movement of a cross point of two Kikuchi lines to calculate the tilt angle and tilt speed of each frame.

### Nanoparticles data collection

A drop (4 μl) of 10-nm gold colloid (BBI Solutions) was applied to a glow-discharged carbon film (Zhongjingkeyi Technology). After a 10-min waiting time, the grid was blotted manually using a filter paper. The Nanoparticles sample was transferred to microscope using a Gatan 626 cryo-holder but without liquid nitrogen cooling. A DE20 camera operating at the movie mode was used to record the sample tilting with a frame rate of 25 frames/s. The values of the camera initial time and exposure time in eTasED were set slightly larger than the true values to ensure the movie contained the whole process of a tilt. Then movies of gold particles moving with stage tilting for 1.4° with 1% speed were collected. These movies were processed using a custom-made MATLAB script to measure the movement of the gold particles.

---

**Supplementary figure 1.**
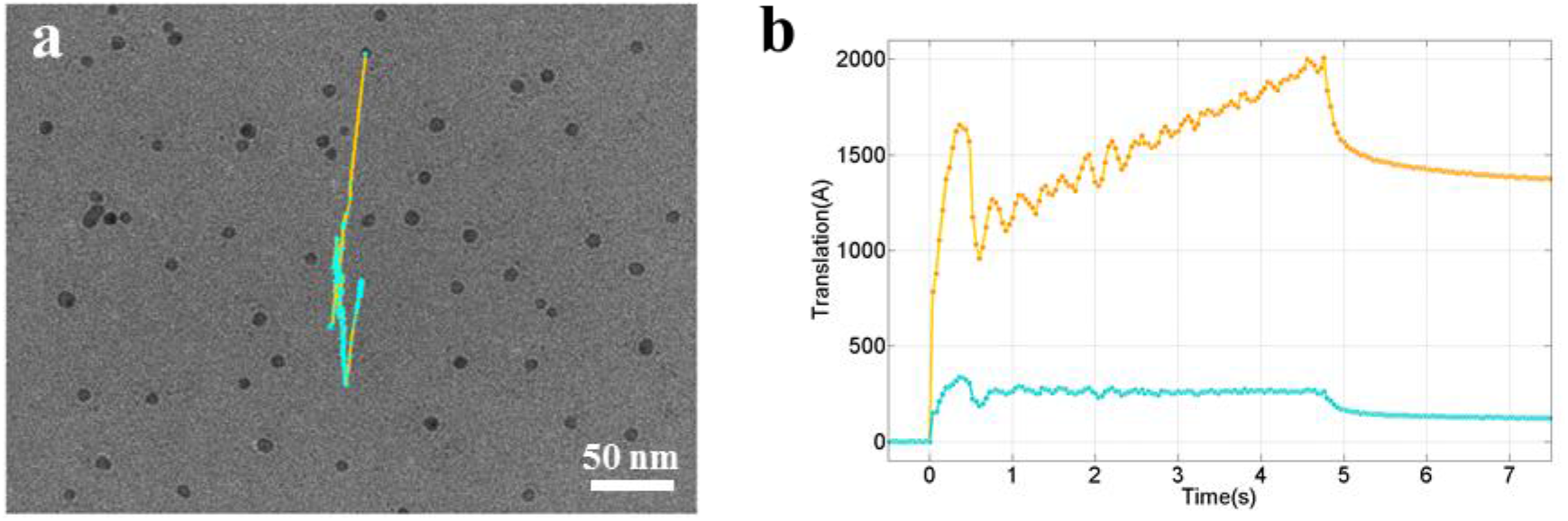
Behavior of CompuStage during continuous tilting. **a)** The trace of the motion during a tilt of 1.4°. The background is an image of nanoparticles observed by F20. A video of the motion of the nanoparticles are shown in Supplementary Movie 2. **b)** The translations of **a)** measured along (orange curve) and perpendicular to (cyan curve) the tilting direction. In the beginning of the tilting, CompuStage indicated a strong and large motion, and then partially recovered. With the tilting, the stage shakes along the tilting direction. The shaking becomes weaker gradually with increasing time. After the tilting stops, the stage moves back slowly. The shaking direction is not exactly along the tilting direction, and has a component perpendicular to the tilting direction.

**Supplementary Figure 2.**
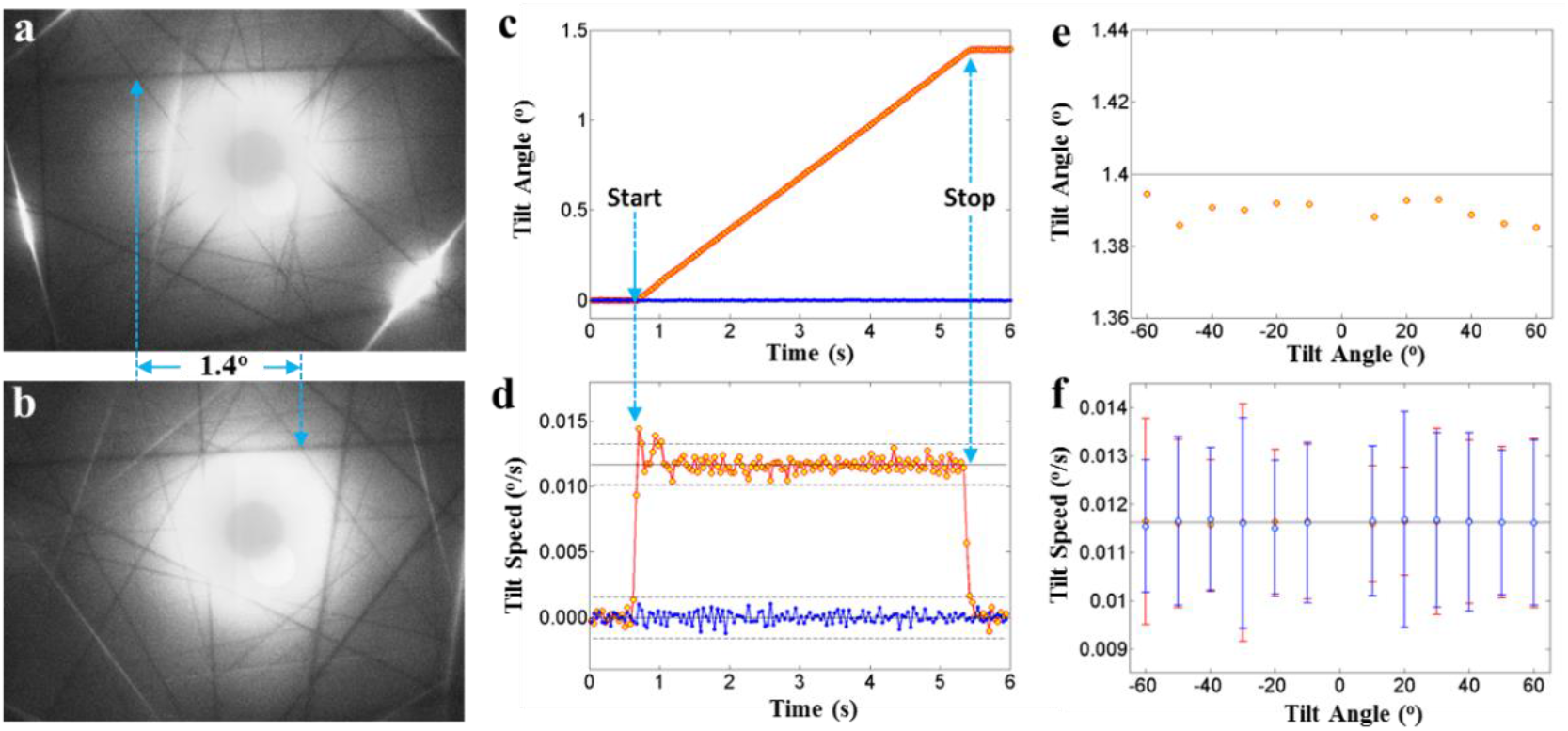
Stability of CompuStage during the continuous tilting. **a)** and **b)** are the Kikuchi patterns before and after a tilt of 1.4° (Supplementary Movie 1), respectively. The cross point of two Kikuchi lines, used for the motion tracking, is indicated by the blue arrows. **c)** and **d)** are the curves of the tilt angle and speed against the time, respectively. The orange curves are for the angle or speed along the tilting direction, and the blue curves are perpendicular to the tilting direction. **e)** Scatter plot of the measured tilt angles of 1.4° against the starting angles. **f)** Scatter plot of the tilt speed against the starting angle. The orange points are for the speed along the tilting direction, and the blue points are for that perpendicular to the tilting direction. Error bars of the tilt speeds are shown correspondingly.

**Supplementary Figure 3.**
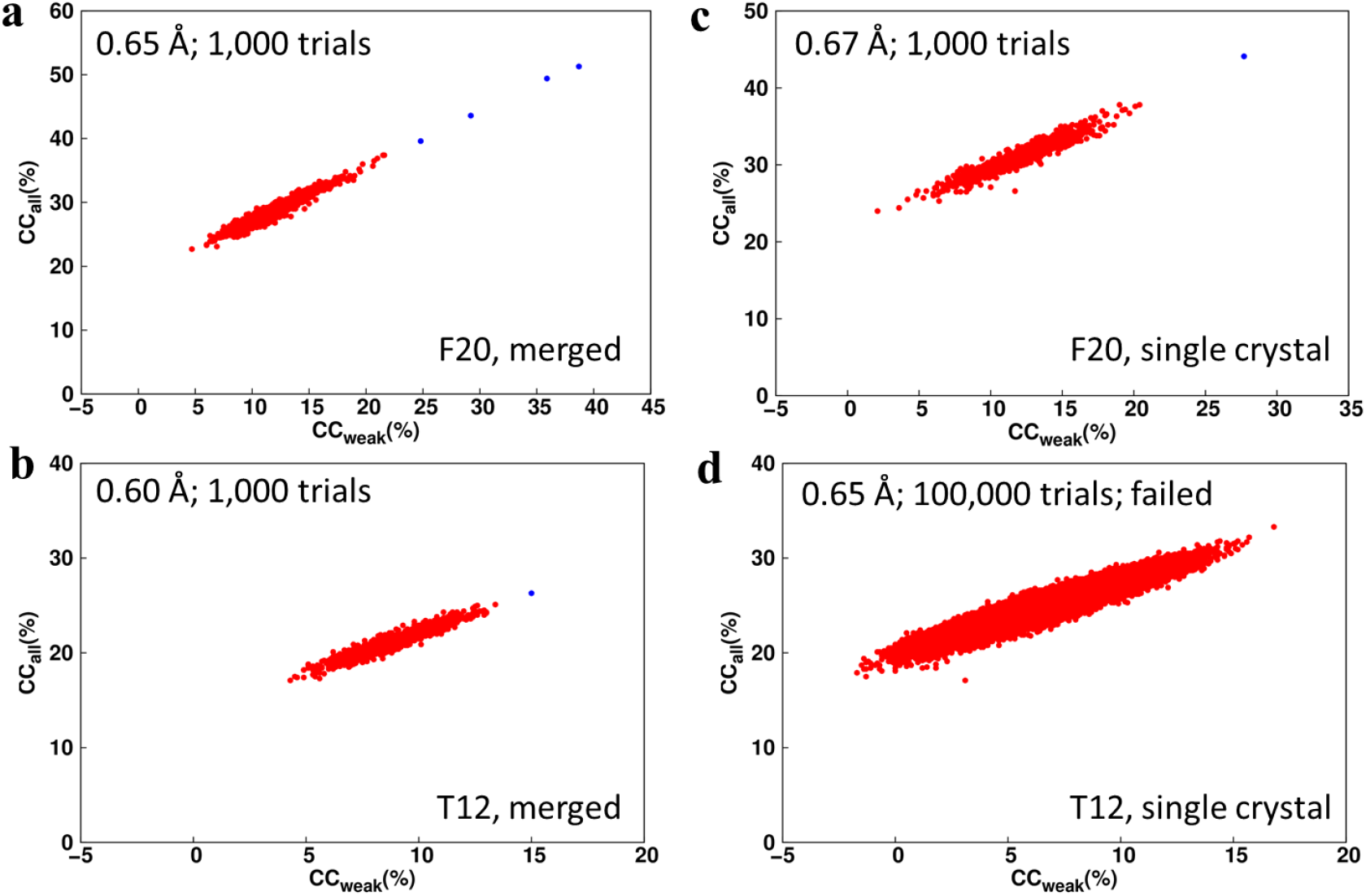
Structure solution of the merged and single-crystal datasets collected on F20 and T12. **a)** and **b)** are the scatter plots of CC_all_ against CC_weak_ of the merged datasets from F20 and T12, respectively. **c)** and **d)** are the scatter plots of CC_all_ against CC_weak_ of the single-crystal datasets from F20 and T12, respectively. Both CC_all_ and CC_weak_ were calculated by SHELXD. The correct solutions correspond to the blue dots, and the unsuccessful trials are shown as red dots. The resolution of each dataset and the number of trials were annotated in each plot.

**Supplementary Figure 4.**
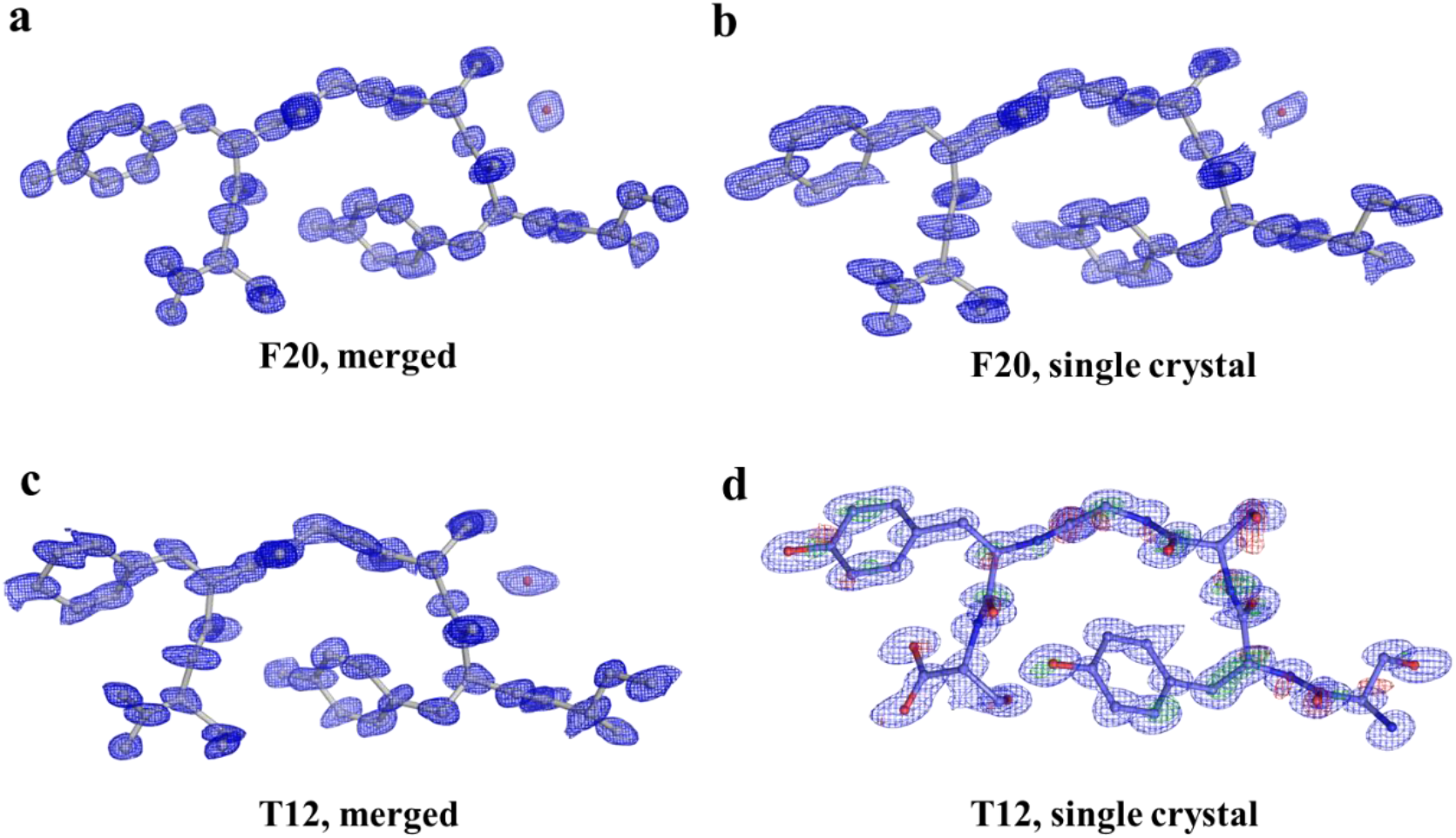
The models and density maps calculated from different datasets. Except the single-crystal T12 dataset that was solved by molecular replacement, all the other three datasets were successfully solved by the direct method. **a)** The merged F20 dataset. **b)** The single-crystal F20 dataset. **c)** The merged T12 dataset. The models are shown in a gray ball-and-stick model, and the density maps are presented by blue mesh. **d)** The single-crystal T12 dataset. The model is presented as ball-and-stick (C, lightblue; O, red; N, blue). The 2F_o_-F_c_ map is drawn as blue mesh with contour threshold at 2.0 σ. The F_o_-F_c_ map is drawn as green (contour threshold at 3.0 σ) and red (contour threshold at −3.0σ) maps. All the models and maps were calculated by SHELXD.

**Supplementary Figure 5.**
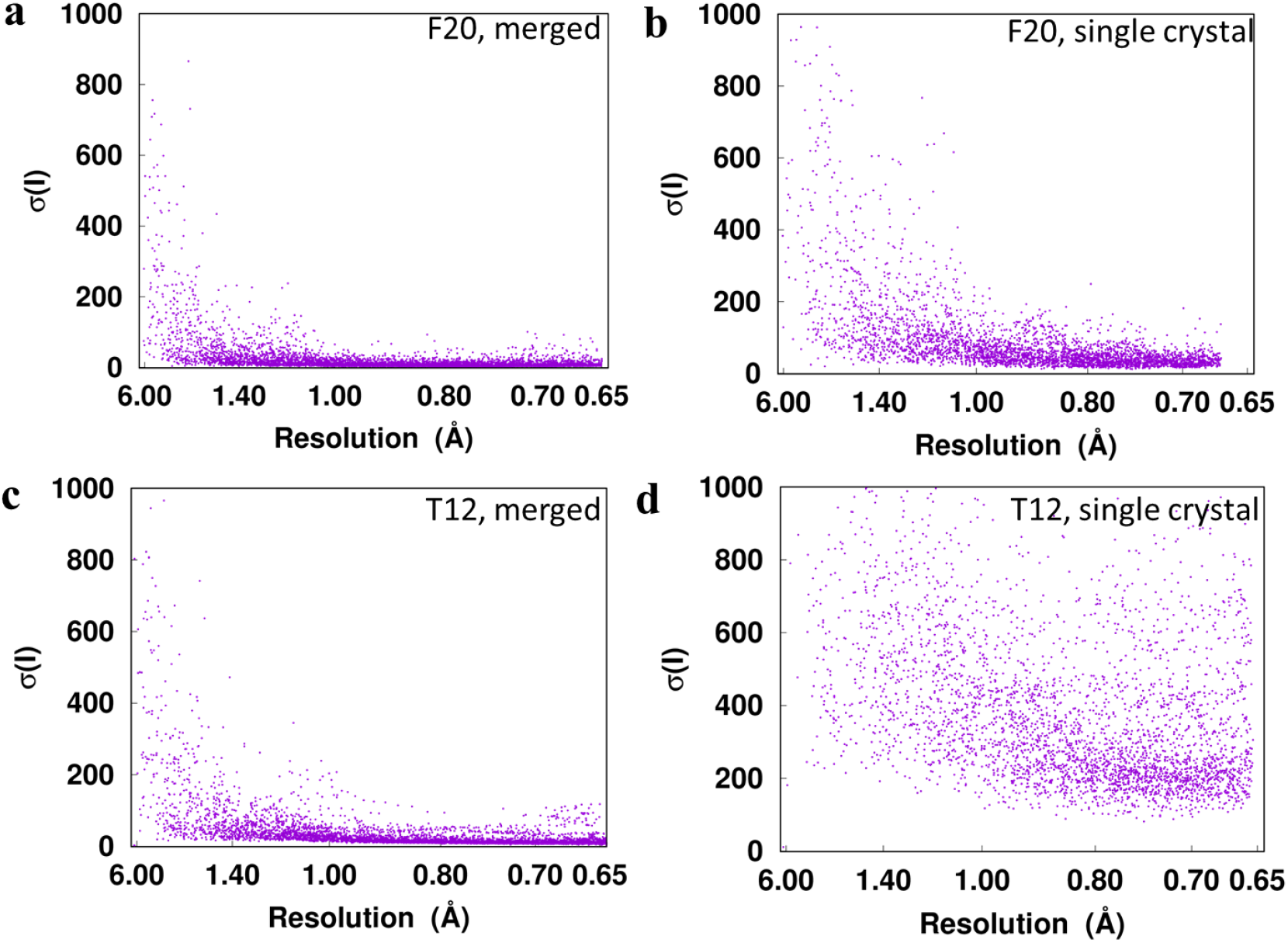
The distribution of σ(I) of the merged and single-crystal datasets. **a)** The merged F20 dataset. **b)** The single-crystal F20 dataset. **c)** The merged T12 dataset. **d)** The single-crystal T12 dataset.

**Supplementary Figure 6.**
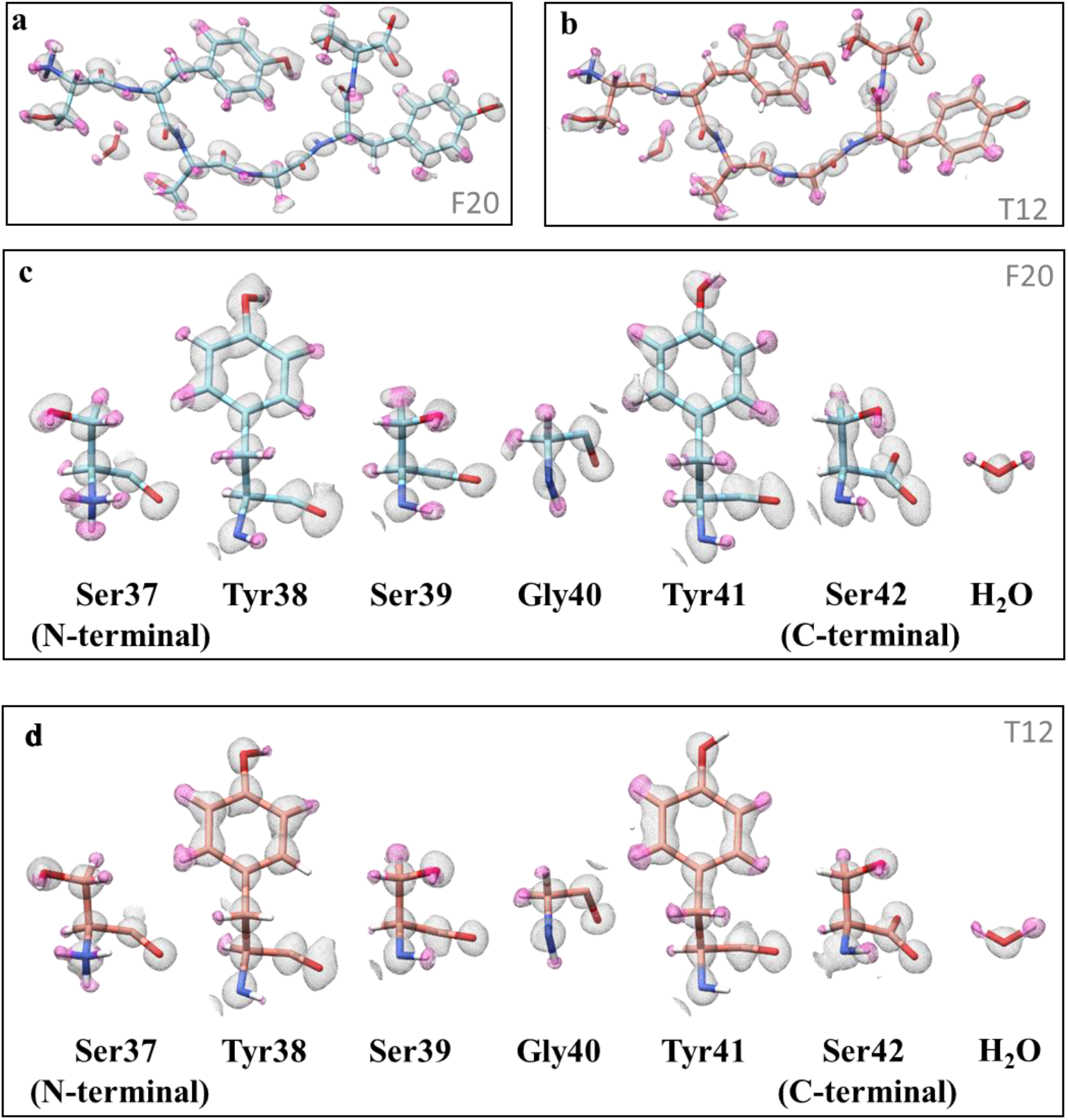
Structures solved from the single-crystal datasets. a) and b) are *2F_o_-F_c_* density maps solved from the single-crystal datasets from F20 and T12, respectively. c) and d) are the magnified density maps of the residues in a) and b), respectively. The densities of the hydrogen atoms are colored in pink. The contour threshold of all the map are at 1.0 σ.

**Supplementary Figure 7.**
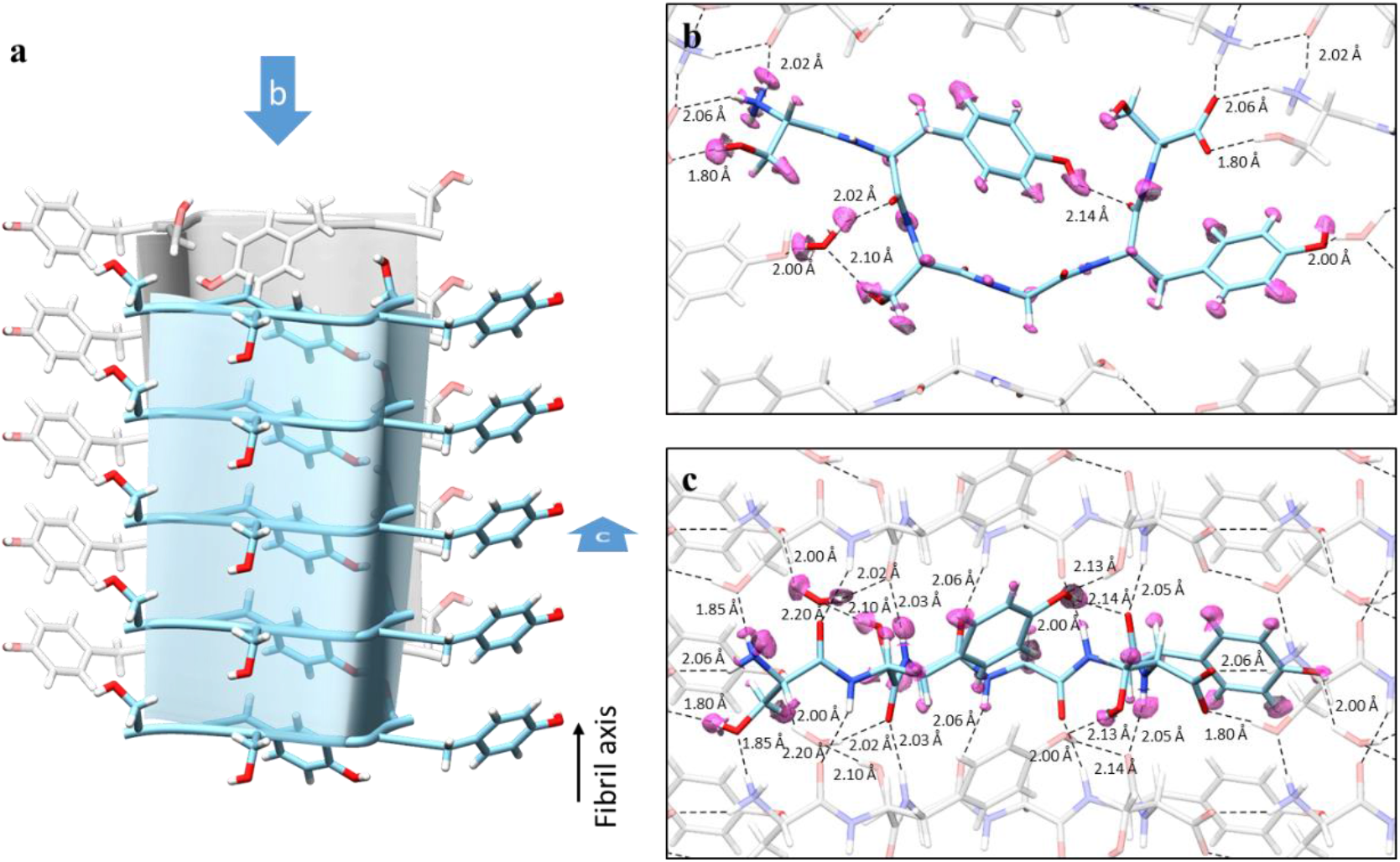
The determined crystal structure of FUS LC RAC1. **a)** The arrangement of the FUS LC RAC1 peptides in crystal. **b)** and **c)** are the hydrogen-bond network shown in two perpendicular views. The density maps of the hydrogen atoms are drawn in pink. The dashed lines are the hydrogen bonds, and the length of the bonds are annotated.

**Supplementary Figure 8.**
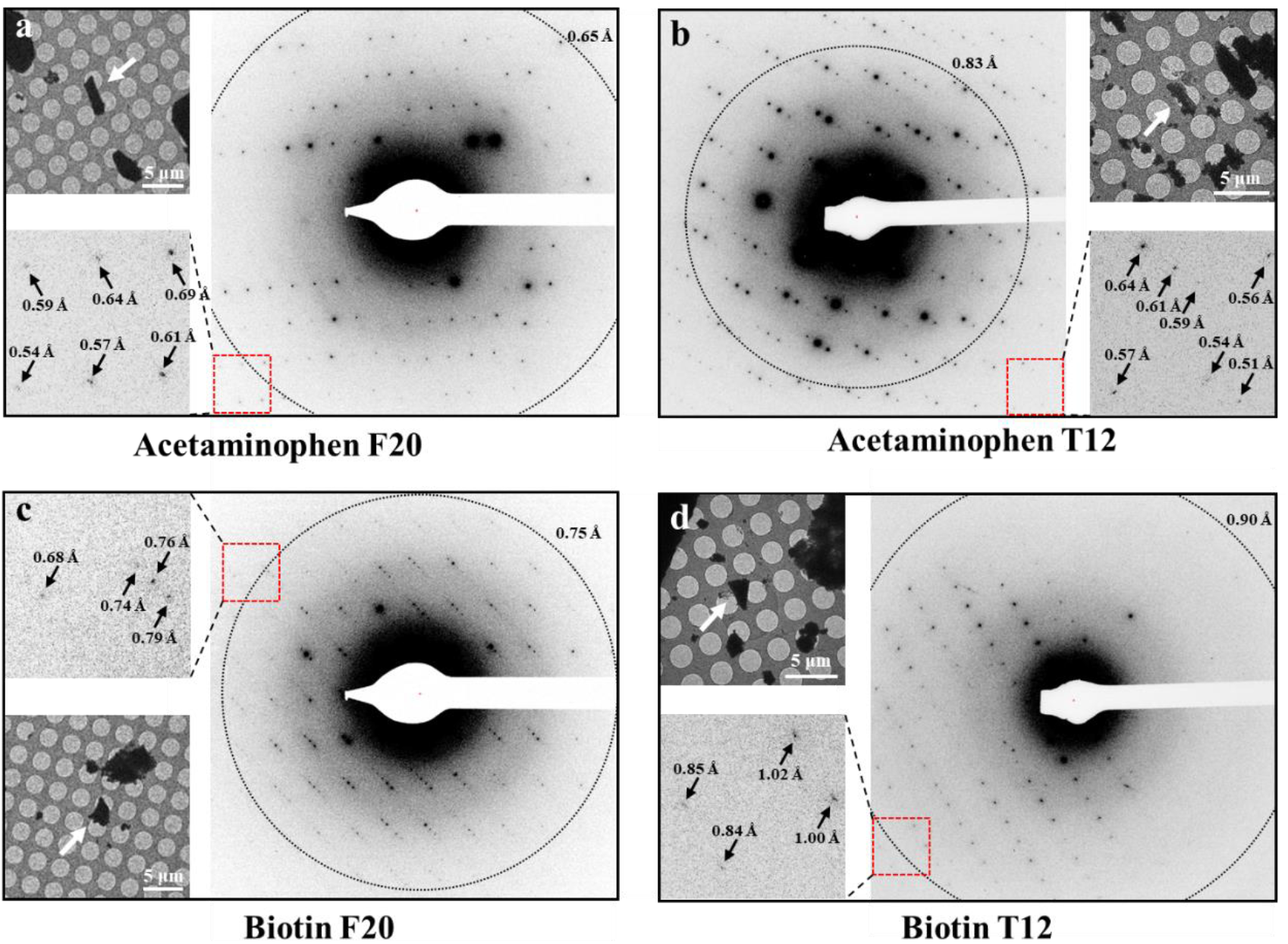
Typical diffraction patterns of Acetaminophen and Biotin crystals. **a)** and **b)** are diffraction patterns of Acetaminophen crystals collected on F20 and T12, respectively. **c)** and **d)** are diffraction patterns of Biotin crystals collected on F20 and T12, respectively. The red crosses in the center of the diffraction patters indicate the beam center, and the dotted rings indicate the final resolution of the solved structures. The arrows on the magnified images of the red boxes indicate some high-resolution diffractions. The corresponding images of the needle-like crystals (pointed by white arrows) are shown on each panel.

**Supplementary Table 1.**
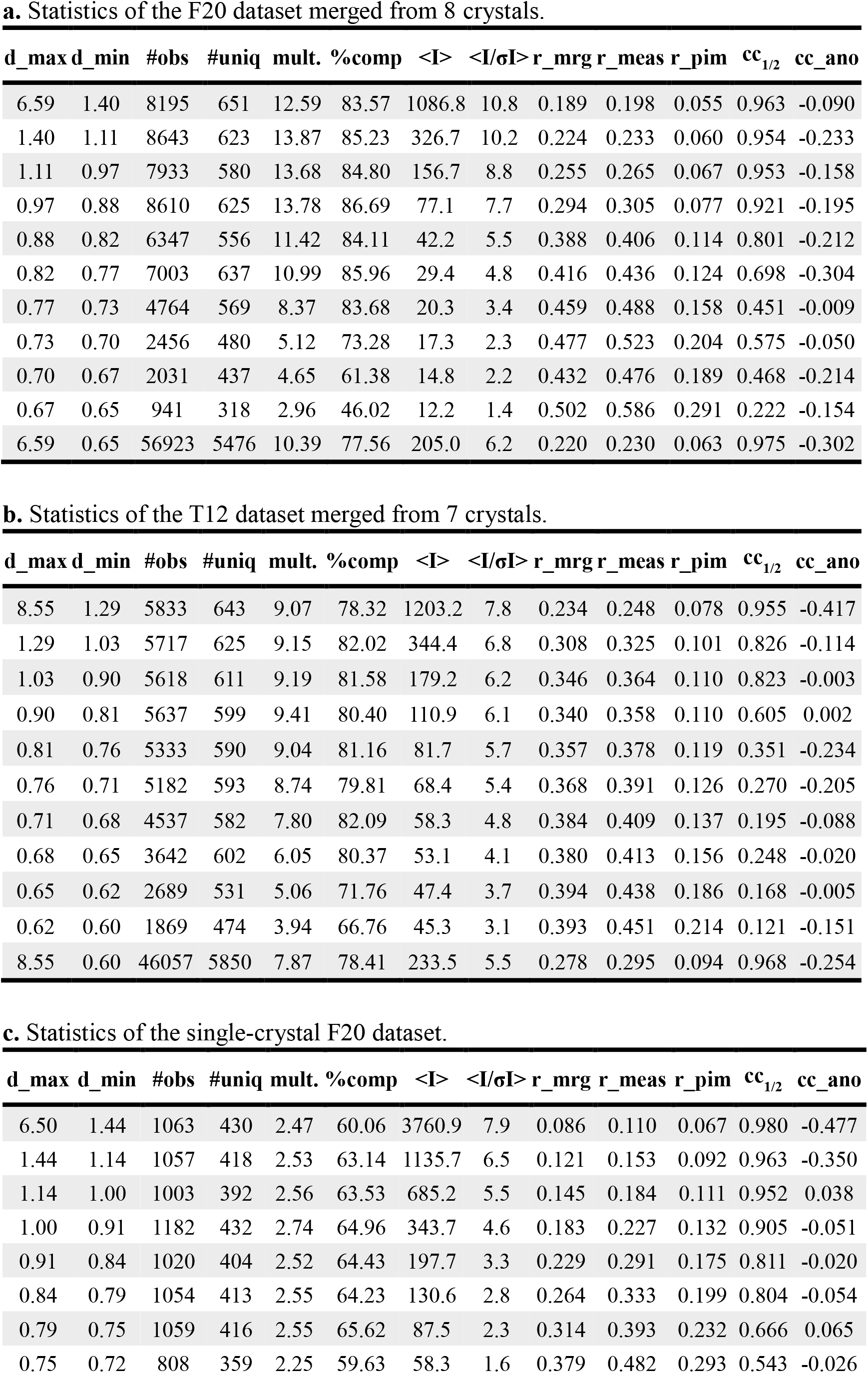
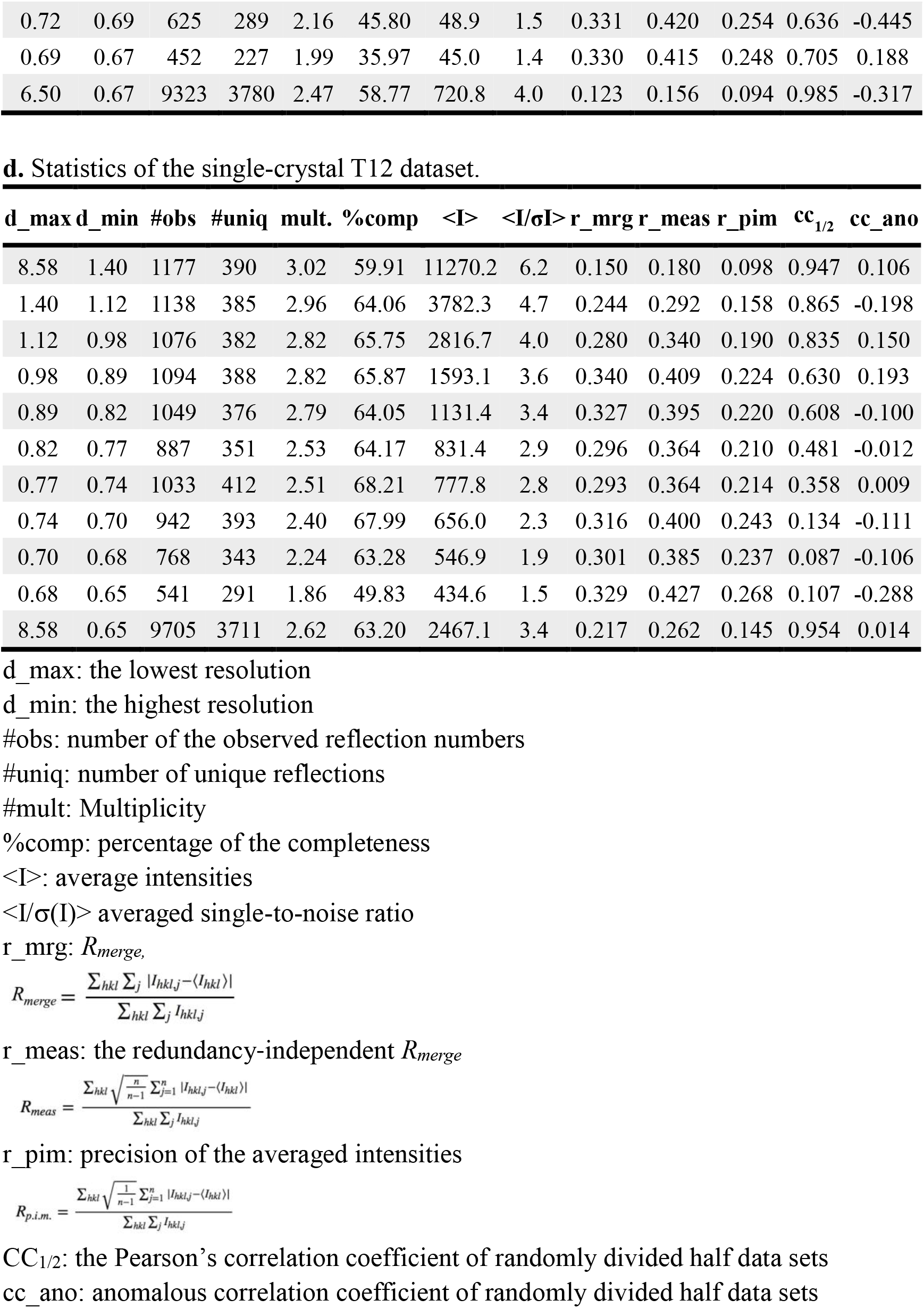
Statistics of the diffraction qualities.

**Supplementary Table 2.**
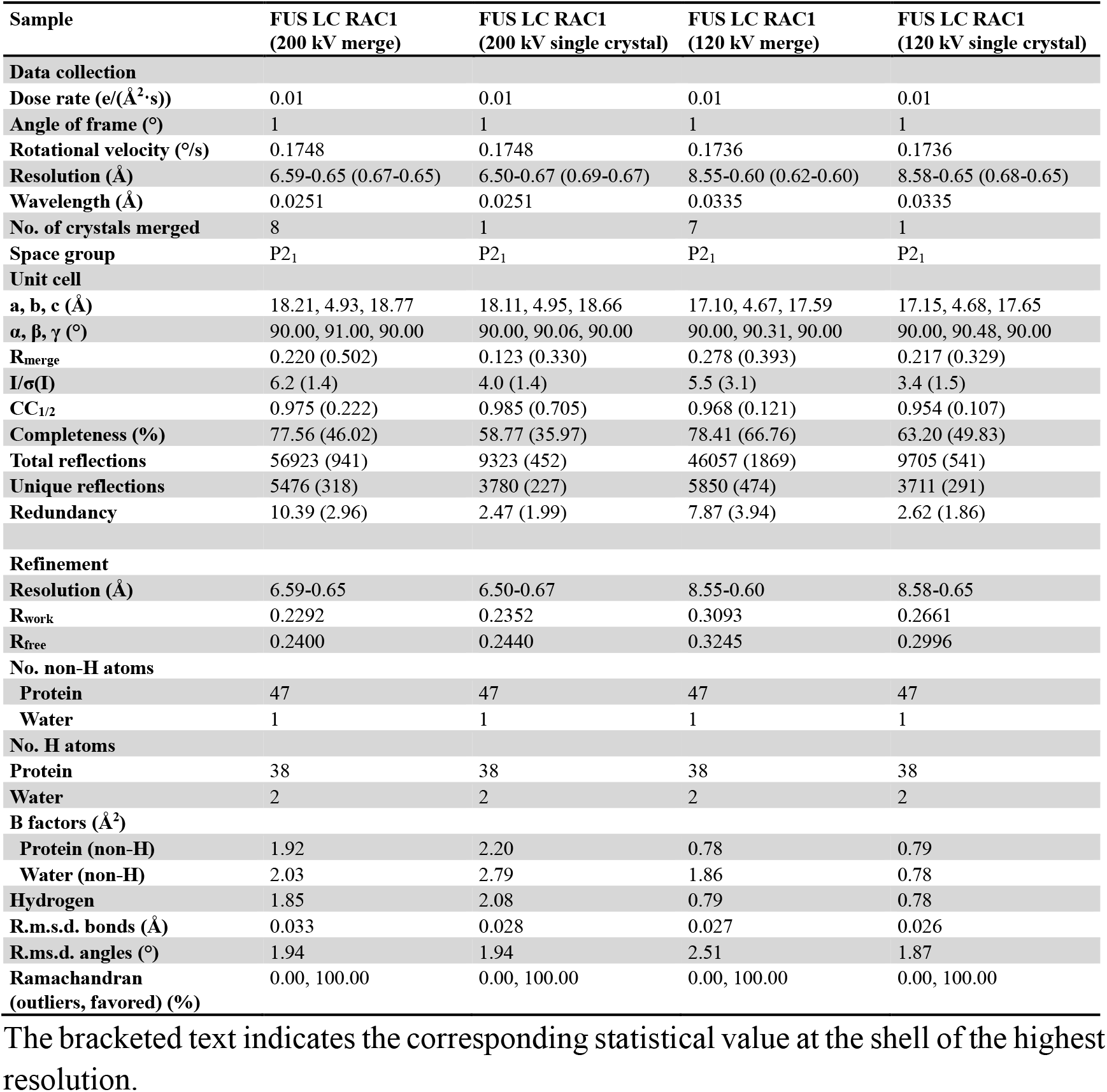
Statistics of the data collection and structure refinement.

| Sample | FUS LC RAC1<br>(200 kV merge) | FUS LC RAC1<br>(200 kV single crystal) | FUS LC RAC1<br>(120 kV merge) | FUS LC RAC1<br>(120 kV single crystal) |
| --- | --- | --- | --- | --- |
| **Data collection** |  |  |  |  |
| Dose rate (e/(Å<sup>2</sup>·s)) | 0.01 | 0.01 | 0.01 | 0.01 |
| Angle of frame (°) | 1 | 1 | 1 | 1 |
| Rotational velocity (°/s) | 0.1748 | 0.1748 | 0.1736 | 0.1736 |
| Resolution (Å) | 6.59-0.65 (0.67-0.65) | 6.50-0.67 (0.69-0.67) | 8.55-0.60 (0.62-0.60) | 8.58-0.65 (0.68-0.65) |
| Wavelength (Å) | 0.0251 | 0.0251 | 0.0335 | 0.0335 |
| No. of crystals merged | 8 | 1 | 7 | 1 |
| Space group | P2<sub>1</sub> | P2<sub>1</sub> | P2<sub>1</sub> | P2<sub>1</sub> |
| **Unit cell** |  |  |  |  |
| a, b, c (Å) | 18.21, 4.93, 18.77 | 18.11, 4.95, 18.66 | 17.10, 4.67, 17.59 | 17.15, 4.68, 17.65 |
| α, β, γ (°) | 90.00, 91.00, 90.00 | 90.00, 90.06, 90.00 | 90.00, 90.31, 90.00 | 90.00, 90.48, 90.00 |
| R<sub>merge</sub> | 0.220 (0.502) | 0.123 (0.330) | 0.278 (0.393) | 0.217 (0.329) |
| I/σ(I) | 6.2 (1.4) | 4.0 (1.4) | 5.5 (3.1) | 3.4 (1.5) |
| CC<sub>1/2</sub> | 0.975 (0.222) | 0.985 (0.705) | 0.968 (0.121) | 0.954 (0.107) |
| Completeness (%) | 77.56 (46.02) | 58.77 (35.97) | 78.41 (66.76) | 63.20 (49.83) |
| Total reflections | 56923 (941) | 9323 (452) | 46057 (1869) | 9705 (541) |
| Unique reflections | 5476 (318) | 3780 (227) | 5850 (474) | 3711 (291) |
| Redundancy | 10.39 (2.96) | 2.47 (1.99) | 7.87 (3.94) | 2.62 (1.86) |
| **Refinement** |  |  |  |  |
| Resolution (Å) | 6.59-0.65 | 6.50-0.67 | 8.55-0.60 | 8.58-0.65 |
| R<sub>work</sub> | 0.2292 | 0.2352 | 0.3093 | 0.2661 |
| R<sub>free</sub> | 0.2400 | 0.2440 | 0.3245 | 0.2996 |
| **No. non-H atoms** |  |  |  |  |
| Protein | 47 | 47 | 47 | 47 |
| Water | 1 | 1 | 1 | 1 |
| **No. H atoms** |  |  |  |  |
| Protein | 38 | 38 | 38 | 38 |
| Water | 2 | 2 | 2 | 2 |
| **B factors (Å<sup>2</sup>)** |  |  |  |  |
| Protein (non-H) | 1.92 | 2.20 | 0.78 | 0.79 |
| Water (non-H) | 2.03 | 2.79 | 1.86 | 0.78 |
| Hydrogen | 1.85 | 2.08 | 0.79 | 0.78 |
| R.m.s.d. bonds (Å) | 0.033 | 0.028 | 0.027 | 0.026 |
| R.m.s.d. angles (°) | 1.94 | 1.94 | 2.51 | 1.87 |
| Ramachandran<br>(outliers, favored) (%) | 0.00, 100.00 | 0.00, 100.00 | 0.00, 100.00 | 0.00, 100.00 |

The bracketed text indicates the corresponding statistical value at the shell of the highest resolution.

**Supplementary Table 3.**
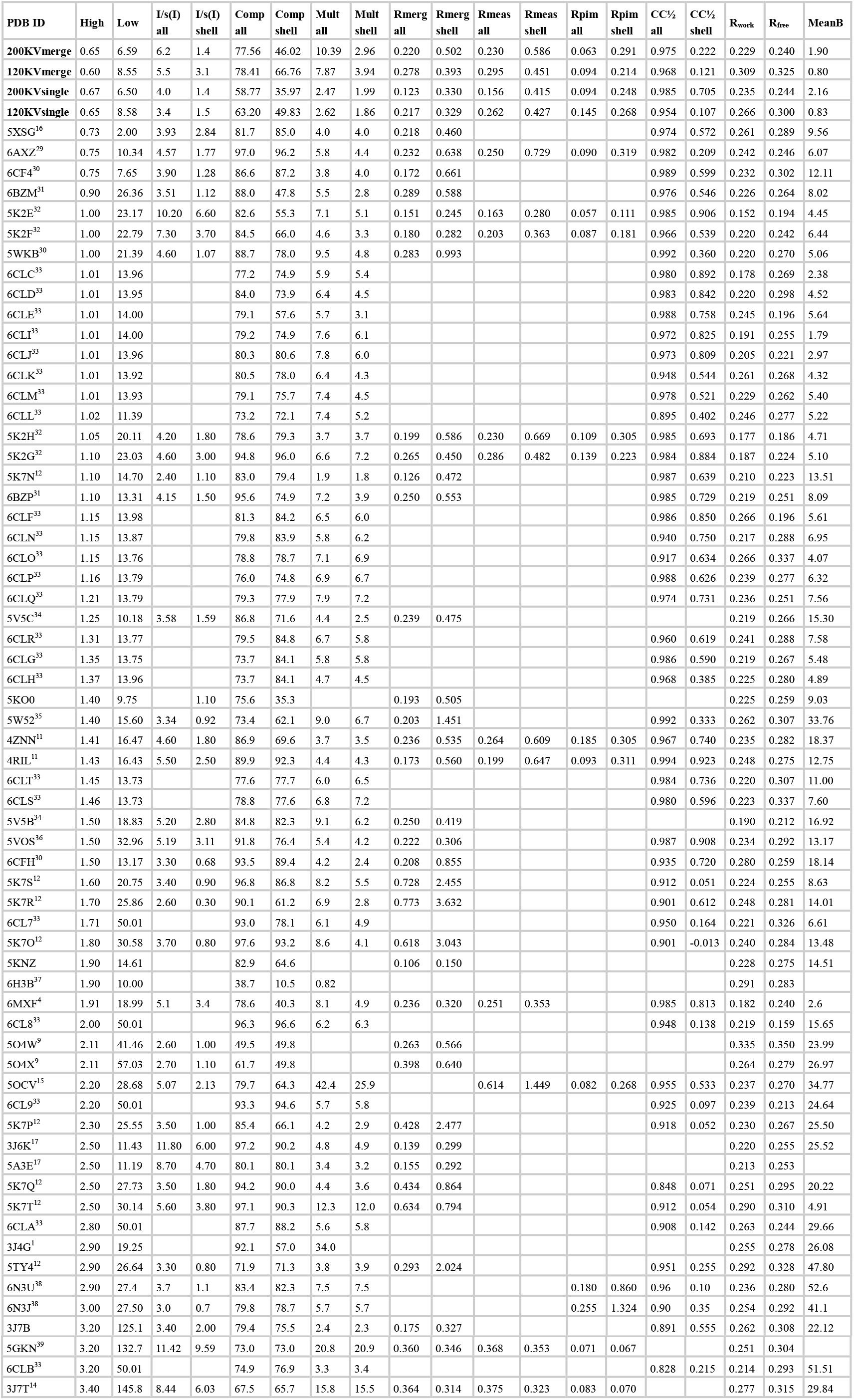
Statistics of all available MicroED data sorted by resolution (Jun. 12, 2018). The first four rows are the results of FUS LC RAC1 crystals in the present work.

**Supplementary Table 4.** Statistics of the data collection and structure refinement of Acetaminophen and Biotin.

|  | Acetaminophen |  | Biotin |  |
| --- | --- | --- | --- | --- |
| Stoichiometric formula | C<sub>8</sub> H<sub>9</sub> N<sub>1</sub> O<sub>2</sub> |  | C<sub>10</sub> H<sub>16</sub> N<sub>2</sub> O<sub>3</sub> S |  |
| Temperature (K) | 100 |  | 100 |  |
| Space group | *P*2<sub>1</sub>/n |  | *P*2<sub>1</sub> 2<sub>1</sub> 2<sub>1</sub> |  |
| Microscope | F20 | T12 | F20 | T12 |
| Number of crystals | 3 | 3 | 4 | 3 |
| Lattice lengths a, b, c (Å) | 7.37, 9.37, 12.09 | 7.06, 9.18, 11.61 | 5.38, 10.77, 21.91 | 5.19, 10.27, 20.88 |
| Lattice angles α, β, γ (°) | 90.00, 98.01, 90.00 | 90.00, 97.22, 90.00 | 90.00, 90.00, 90.00 | 90.00, 90.00, 90.00 |
| Num of Reflections | 11760 (485) | 2569 (54) | 11177 (1033) | 3672 (183) |
| Num of Unique reflections | 3038 (200) | 948 (44) | 1838 (178) | 890 (76) |
| *R*<sub>obs</sub> | 0.248 (0.375) | 0.182 (0.291) | 0.248 (0.765) | 0.252 (0.332) |
| *R*<sub>meas</sub> | 0.282 (0.454) | 0.219 (0.369) | 0.273 (0.834) | 0.286 (0.418) |
| Multiplicity | 3.87 (2.43) | 2.71 (1.16) | 6.08 (5.80) | 4.13 (2.41) |
| *I*/σ(*I*) | 3.0 (1.7) | 3.7 (2.0) | 3.4 (1.7) | 3.8 (2.0) |
| *CC*<sub>1/2</sub> (%) | 91.4 (73.0) | 94.5(69.0) | 97.3 (42.6) | 92.9(21.2) |
| Resolution (Å) | 0.65 | 0.83 | 0.75 | 0.90 |
| Completeness (%) | 91.56 (58.82) | 70.96 (34.11) | 99.7 (98.9) | 92.32 (81.72) |
| *R* | 0.242 | 0.224 | 0.266 | 0.189 |
| *wR*<sub>2</sub> | 0.584 | 0.558 | 0.538 | 0.432 |
| *Goof* | 1.215 | 1.289 | 1.679 | 1.264 |

**Supplementary Movie 1.** The movement of gold nano-particles under continuous tilting of the stage.

**Supplementary Movie 2.** The movement of Kikuchi lines under continuous tilting of the stage.

